# Microscale visualization of cellular features in adult macaque visual cortex

**DOI:** 10.1101/2023.11.02.565381

**Authors:** Pooja Balaram, Kevin Takasaki, Ayana Hellevik, Jamuna Tandukar, Emily Turschak, Bryan MacLennan, Naveen Ouellette, Russel Torres, Connor Laughland, Olga Gliko, Sharmistaa Seshamani, Eric Perlman, Mike Taormina, Erica Peterson, Zoe Juneau, Lydia Potekhina, Adam Glaser, Jayaram Chandrashekar, Molly Logsdon, Kevin Cao, Celeste Dylla, Gaku Hatanaka, Soumya Chatterjee, Jonathan Ting, David Vumbaco, Jack Waters, Wyeth Bair, Doris Tsao, Ruixuan Gao, Clay Reid

## Abstract

Expansion microscopy and light sheet imaging enable fine-scale resolution of intracellular features that comprise neural circuits. Most current techniques visualize sparsely distributed features across whole brains or densely distributed features within individual brain regions. Here, we visualize dense distributions of immunolabeled proteins across early visual cortical areas in adult macaque monkeys. This process may be combined with multiphoton or magnetic resonance imaging to produce multimodal atlases in large, gyrencephalic brains.

## Main Text

Expansion microscopy (ExM) facilitates nanoscale imaging of biological structures with diffraction-limited microscopy in a wide variety of tissue preparations.^1–13^ In brain tissue, ExM enables discrete visualization of hydrophilic biomolecules such as proteins, DNA, and mRNA, which are densely packed within and around hydrophobic myelin sheaths, by physically separating hydrophobic and hydrophilic tissue regions through isotropic expansion ^3,7,12,13^. Detailed analyses of cellular features across large samples of brain tissue become feasible when ExM is combined with light sheet fluorescence microscopy (LSFM), which accelerates acquisition of high resolution image data in cleared tissues.^14–17^ This combinatorial technique is particularly useful when investigating neuroanatomy and connectivity in large, gyrencephalic nonhuman primate and human brains,^5,11–14^ where prior studies are limited to sparsely labeled neural populations or subsets of cellular features within neural circuits.^18^

Here, we adapt standard protocols for proExM^3,6^ and LSFM^19^ to visualize intracellular features in densely packed neuronal populations across primary (V1) and secondary (V2) visual cortex of adult macaque monkeys. V1 and V2 span a significant portion of both cortical occipital lobes in macaques, and neurons in these areas are organized into discrete functional domains that streamline processing of color, form and motion in complex visual scenes.^20–24^ Subsets of feedforward projections from V1 have been previously identified via immunolabeling for non-phosphorylated heavy-chain neurofilaments,^25–30^ but the morphology and connectivity of individual neurons within these circuits is largely unknown. Our protocols for ExM and LSFM in immunolabeled tissue from macaque visual cortex (Figure 1) enable visualization of individual neurons and efferent projections within anatomically defined visual circuits.

**Figure 1.**
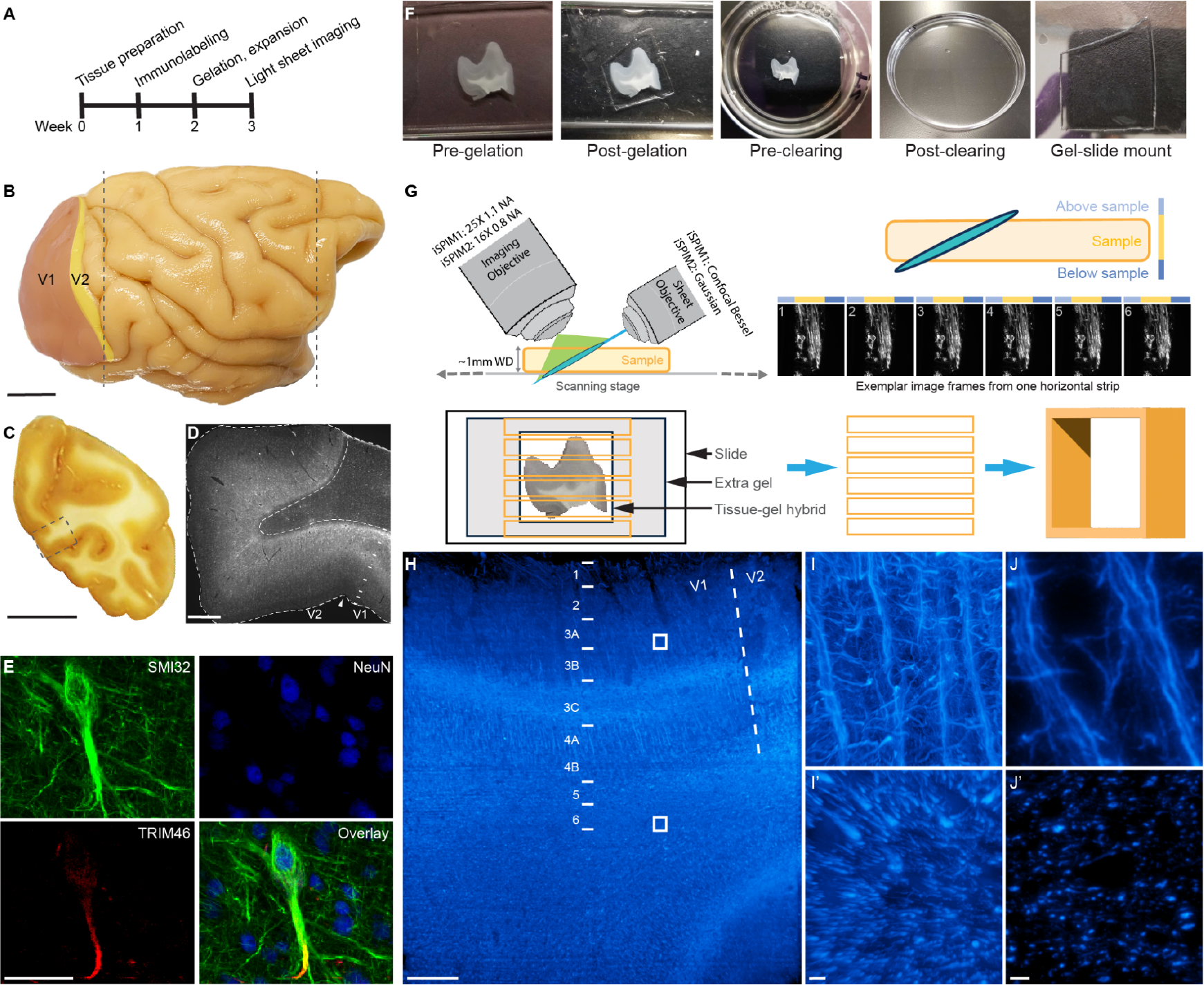
Microscale visualization of densely distributed cellular features in visual cortex of macaque monkeys. (A) Experimental timeline. (B) Cortical hemispheres from adult macaque monkeys were extracted following transcardial perfusion and blocked along the rostral and caudal poles of the temporal lobe (gray dotted lines). Caudal blocks containing primary (V1) and secondary (V2) visual cortical areas were then cryosectioned coronally at 100 *μ*m. Scale bar is 1 cm. (C) In each cryosection, tissue regions containing the V1/V2 border along the lunate, calcarine, or inferior occipital sulcus were manually dissected prior to immunolabeling. Gray dotted outline marks location of tissue regions depicted in panel D. Scale bar is 1 cm. (D) Immunolabeled, expanded tissue-hydrogel containing the V1/V2 border along the upper bank of the calcarine sulcus. Boundaries between gray and white matter regions (dotted lines), visual cortical areas (arrowhead) and intracortical layers (solid lines) are clearly visible based on changes in fluorescent labeling intensity post-expansion. Scale bar is 1 mm. (E) Antibodies against non-phosphorylated heavy-chain neurofilaments (SMI32), neuronal cell bodies (NeuN), and axon initial segments (TRIM46), can be combined in individual tissue sections to visualize neuronal morphology and local connectivity. Scale bar is 50 *μ*m. (F) Custom ExM protocols, including modifications to linker concentrations and digestion conditions, produced transparent tissue-hydrogel matrices with ∼4X expansion in hydrophobic gray matter and hydrophilic white matter regions of visual cortex. Labeled, cleared, and expanded hydrogels were then mounted on poly-L-lysine coated coverslips prior to imaging (G) Hydrogels were light sheet imaged at 16X or 25X using custom inverted selective plane illumination microscopes (iSPIMs). Each hydrogel was imaged in horizontal blocks; image blocks were then concatenated to produce 3D volumes of tissue regions of interest within each hydrogel. (H) Exaspim imaging of an immunolabeled, expanded hydrogel spanning the V1/V2 border along the lower bank of the calcarine sulcus. Cortical areal boundaries (dotted line) and intracortical layers (solid lines) can be defined based on changes in SMI32 labeling intensity. Boxes demarcate tissue regions of interest in cortical gray matter and subcortical white matter depicted in subsequent panels. Scale bar is 1 mm. (I, I’) Maximum intensity projections of SMI32 immunolabeling spanning the full depth of the EXAspim volume identify unique patterns of non-phosphorylated heavy-chain neurofilaments in superficial cortical layers (I) and deep white matter (I’) in V1. Scale bar is 50 *μ*m. (J, J’) Single-plane visualizations of SMI32 immunolabeling through the middle of the EXAspim volume demonstrate discrete voxels of fluorescent label and negligent background labeling, which enables high ratios of signal-to-noise in large image volumes. Scale bar is 10 *μ*m.

First, we streamlined sample preparation to minimize variations in tissue quality. Transcardially perfused cortical hemispheres were blocked at the rostral and caudal poles of the temporal lobe, such that early visual cortical areas in each occipital lobe were contained in a single block (Fig 1B). Each occipital block was serially sectioned on a sliding freezing microtome to minimize microscale distortions at the boundaries of adjacent sections, and tissue regions along the calcarine, lunate, or inferior occipital sulcus containing V1 and V2 were manually dissected from each section (Fig. 1C). Gross morphology, gray and white matter boundaries, and areal and laminar boundaries were well preserved in each sample following immunolabeling, clearing and expansion (Fig. 1D), which facilitates registration of expanded hydrogels to the original sample.

Next, we developed a fluorescent immunolabeling strategy that demarcated neuronal cell bodies, axon initial segments, and non-phosphorylated heavy-chain neurofilaments in individual sections; when combined, these labels visualize single neurons and their efferent projections in layers 3C and 5 of V1 (Figs. 1E, S2).^25,28^ Immunolabeled sections were then incubated with high concentrations of 6-((acryloyl)amino)hexanoic acid (acryloyl-X or AcX) in acidified phosphate-buffered saline to increase AcX penetration through densely myelinated tissue regions and maximize anchoring of fluorescent secondary antibodies to the hydrogel polymer network. Expandable hydrogels were then formed using previously described ratios of sodium acrylate to acrylamide^6^ to achieve ∼4X expansion of each sample when immersed in distilled deionized water (Fig. 1D, F). Unbound tissue components were removed from each hydrogel through a combination of enzymatic digestion with Proteinase K and mechanical homogenization with sodium dodecyl sulfate and Triton-X-100,^4^ which produced consistent clearing in both hydrophilic gray matter and hydrophobic white matter tissue regions (Fig. 1E). Once expanded in water, each hydrogel was mounted on a poly-L-lysine coated coverslip as previously described.^6^

Mounted hydrogels were imaged on custom selective plane illumination microscopes fitted with commercial scanning stages and 16X 0.8 NA or 25X 1.1 NA objectives (Fig. 1G). Image data was acquired in horizontal strips along the longest axis of each sample at resolutions ranging from 1 to 5 *μ*m/pixel. ‘Overview’ images of each hydrogel were first acquired to locate tissue regions of interest (ROIs) within each sample, followed by full-resolution imaging of each ROI. Following this paradigm, a cleared, expanded 1cm x 1cm x 500*μ*m sample could be imaged in ∼30 minutes. Image strips were then cropped to remove non-tissue regions, deskewed to correct for the angle of acquisition, and concatenated in 2D and 3D for image visualization with commercial and open-source software (Fiji, BigStitcher; Imaris; Neuroglancer).

Combining proExM and LSFM in 100 *μ*m-thick sections of macaque cortex, particularly when visualizing dense distributions of immunolabeled proteins, represents a significant technical advantage over traditional histological preparations and imaging methods. Ultrastructural features such as cytoskeletal neurofilament matrices, previously visualized with confocal or electron microscopic techniques,^31–33^ can be imaged with diffraction-limited light microscopes when cleared and expanded via proExM. Moreover, terabyte quantities of microscale resolution image data spanning centimeters of expanded tissue can be acquired in hours instead of days; a significant improvement over conventional imaging techniques such as confocal microscopy. However, despite these advantages, the collection and subsequent registration of tens to hundreds of serial sections spanning one occipital lobe becomes a challenging and time-consuming process when working with large nonhuman primate brains. To further accelerate data acquisition speeds, we utilized the ExaSPIM imaging system^14^ to visualize immunolabeled neurofilaments across centimeters of a labeled, cleared and expanded hydrogel (Figs 1H-J’).

To expand technical applications of proExM and LSFM in macaque cortex, we utilized the protocols described above to visualize virally-expressed GCaMP6s and immunolabeled SMI32 in V1 and V2 of adult macaque monkeys that underwent multiphoton imaging prior to brain extraction (Supplementary Fig. 1). In naive tissue sections (Fig. S1C, E, F), GCaMP6s expression appears concentrated around injection sites and variably expressed between gray and white matter regions, which poses a challenge for registration of functionally characterized neurons in superficial cortical layers. However, following clearing and expansion (Fig. S1 G-L), GCaMP6s expression is clearly visible in neuronal somas across cortical layers, enabling registration of functionally characterized neurons within each sample. SMI32 immunolabeling, when combined with viral GCaMP6s expression, can subsequently visualize somatodendritic morphologies and putative projections of GCaMP6s-expressing cells in layers 3C and 5 of V1 (Fig. S1J, L).^27,28^

We also combined proExM and LSFM with functional and diffusion magnetic resonance imaging (MRI) to register macroscale (millimeters/pixel) measures of cortical areal boundaries and white matter tracts with microscale (nanometers/pixel) measures of the same features (Supplementary Fig. 2). Retinotopic maps spanning V1 and V2 were acquired with in vivo 3T fMRI (Fig. S2A, B) and diffusion maps of white matter tracts below V1 and V2 were acquired with ex vivo 3T dMRI (Fig. S2C, D), followed by brain extraction and sample preparation as described above. Four fluorescent labels were combined in each tissue section, visualizing cell nuclei (DAPI) myelinated white matter tracts (MBP), non-phosphorylated heavy-chain neurofilaments (SMI32) and parvalbumin-expressing inhibitory cortical neurons (PV), to demarcate unique areal and laminar distributions across occipital visual areas (Fig. S2E), particularly the boundary between V1 and V2 (Fig. S2F).

The same boundaries can be visualized based on changes in fluorescent intensity in cleared, expanded hydrogels (Fig. S2G, H), which facilitates registration of all four modalities in each experiment.

Investigating the architecture of neural circuits in nonhuman primate brains is historically a challenging pursuit, given the technical challenges of processing tissue from gyrencephalic brains and the time commitment required for imaging and quantifying microscopic features across large tissue samples. Our adaptation of proExM and LSFM significantly reduces those hurdles, and offers potential for future studies characterizing the structure and function of discrete neural circuits across multiple brain areas and experimental modalities.

## Acknowledgements

We thank the following Allen Institute teams for their varied contributions to this project: Animal Care, Transgenic Colony Management, Lab Animal Services, Tissue Processing, Viral Technology, Imaging, and Project Management. We also thank the Washington National Primate Research Center and the California Institute of Technology for veterinary and technical support. This work was funded by NIH; RF1MH117820-01A1 and UG3MH126864-01 to R.C.R.

## Methods

Visual cortical tissue was obtained from three adult macaque monkeys (*Macaca nemestrina)* through the Washington National Primate Center (WaNPRC) at the University of Washington (UW) and one adult macaque monkey (*Macaca mulatta*) through the Tsao laboratory at the University of California, Berkeley. All experiments were approved by Institutional Animal Care and Use Committees and followed guidelines established by the National Institutes of Health.

### Sample preparation

Subjects were transcardially perfused with 0.9% sterile saline, followed by 4% paraformaldehyde in saline. Brains were extracted whole, bisected through the midline, and separated into three coronal blocks along the rostral and caudal poles of the temporal lobe. Caudal blocks containing visual cortical areas were cryoprotected in 30% sucrose, sectioned into 100*μ*m coronal sections using a sliding freezing microtome, and stored in cryoprotectant (30% glycerol, 30% ethylene glycol, 40% 1M phosphate buffered saline) at -20°C until further processing.

### Immunolabeling

Free-floating sections containing early visual areas V1-V4 were immunolabeled with SMI32-488 to visualize heavy-chain neurofilament proteins; reagents and conditions are described in Table 1.

**Table 1.** Immunolabeling protocol.

| Phase | Reagents | Temperature, state | Time |
| --- | --- | --- | --- |
| Denaturation | 4M urea in 1M PBS | 4C, orbital shaker | 4 days |
| Blocking solution | 5% normal goat serum, 0.5% Triton, 1M PBS | 4C, orbital shaker | 1 day |
| Antibody conjugate | 1:10 NFH-488 | 4C, orbital shaker | 5 days |

### Hydrogel formation and expansion

Immunolabeled sections were embedded in a hydrogel matrix enabling ∼4X expansion under conditions described in Table 2. Fluorescently labeled proteins were anchored with 6-(acryloyl(amino)) hexanoic acid in acidified phosphate-buffered saline and crosslinked to an expandable polyacrylamide gel. Unanchored tissue components were enzymatically digested with Proteinase K (ProK), with the addition of Triton X-100 and sodium dodecyl sulfate to increase ProK penetration through densely myelinated brain regions. Digested, cleared hydrogels were expanded via serial incubation in a graded series of phosphate-buffered saline to distilled, deionized water (ddH_2_O); complete expansion in each gel was achieved following overnight incubation in ddH_2_O.

Expanded hydrogels were mounted on 2 x 3 x 0.15 mm glass coverslips (Claritex, prior to iSPIM imaging. Each coverslip was briefly dipped in Poly-L-lysine (Millipore Sigma, Rockville MD), air dried, and secured to a flat surface for mounting. Expanded gels were transferred from water onto a non-coated coverslip, dried with Kimwipes or absorbent points to minimize excess liquid along the gel-coverslip interface, and gradually transferred onto the poly-L-lysine coated coverslip. Mounted gels were air dried for 3-5 minutes and stored in ddH_2_0 at room temperature prior to imaging.

### Light sheet imaging and analysis

Light sheet imaging was performed on custom inverted selective plane illumination microscopes (iSPIMs) fit with commercial motorized stages (Applied Scientific Instrumentation, Eugene OR) and objectives (16X 0.8NA 3mm WD, Nikon; 25X 1.1NA 2mm WD, Nikon). Gel coverslips were secured to a stage insert with Kwik-cast (World Precision Instruments, Sarasota FL) and submerged in ddH_2_O for imaging.

Location, orientation, and labeling quality of tissue regions within each gel was assessed by capturing fast-step or ‘overview’ scans (25 *μ*m X step, 1 mm Y step, 800 *μ*m Z step). Overview scans were converted to BigDataViewer (BDV) format and visualized with BigStitcher in Fiji. Regions of interest containing visual cortical areas were then imaged at higher resolution (1 *μ*m X spacing x 700 *μ*m Y spacing) with confocal slit detection to seamlessly capture neuronal features across each ROI. High resolution image data was rendered in Neuroglancer, which visualized labeled features at multiple scales in each tissue volume.

Image regions containing relevant neuronal features such as cell bodies and axons were extracted as separate image volume. Features were semi-automatically segmented using published algorithms for sparse-labeled datasets (ref. IVSCC paper) that were further optimized for densely-labeled datasets. Segmented objects were then overlaid on image data and rendered in Neuroglancer for visualization.

### GCaMP6s labeling of neuronal somas combined with SMI32 labeling of axonal scaffolds

Virally mediated GCaMP6s expression in macaque V1 was achieved with a series of stereotaxic injections of adeno-associated viruses (AAVs) PHP.eB-CAG-GCaMP6s or PHP.eB-hSyn-GCaMP6s spanning the V1/V2 border along the dorsal lunate gyrus. AAVs were generated by the Allen Institute Viral Technology team using commercial plasmids pAAV-CAG-GCaMP6s-WPRE-SV40 and pAAV-Syn-GCaMP6s-WPRE-SV40, both available from Addgene (Plasmids #100844 and #100843 respectively; Addgene, Watertown, MA) courtesy of Dr. Douglas Kim and the GENIE project^34^. Multiphoton imaging of GCaMP6s-expressing cells occurred two to five weeks post-injection, followed by transcardial perfusion with 4% paraformaldehyde. Perfused brain tissue from labeled regions of V1 were cryosectioned, immunolabeled for SMI32, and formed into hydrogels as described above.

### Functional and diffusion magnetic resonance imaging combined with SMI32 labeling of axonal scaffolds

One macaque monkey underwent in vivo 3-T magnetic resonance imaging (fMRI) using standard techniques ^35–37^ prior to transcardial perfusion with 4% paraformaldehyde as described above. Ex-vivo diffusion magnetic resonance imaging was performed using standard techniques ^36^ prior to removing cerebral hemispheres from the skull cavity; once imaging was complete, perfused brain tissue was extracted, blocked and sectioned coronally as described above. One series of coronal sections with ∼500 *μ*m spacing between sections was fluorescently labeled to visualize cell nuclei (DAPI), non-phosphorylated heavy-chain neurofilaments (SMI32), parvalbumin-positive cortical interneurons (PV), and myelin basic protein (MBP). A second series with ∼500 *μ*m spacing, interleaved with the first series, was immunolabeled for SMI32, gelled, cleared, expanded, and light sheet imaged as described above.

## SUPPLEMENTAL INFORMATION

### Author contributions

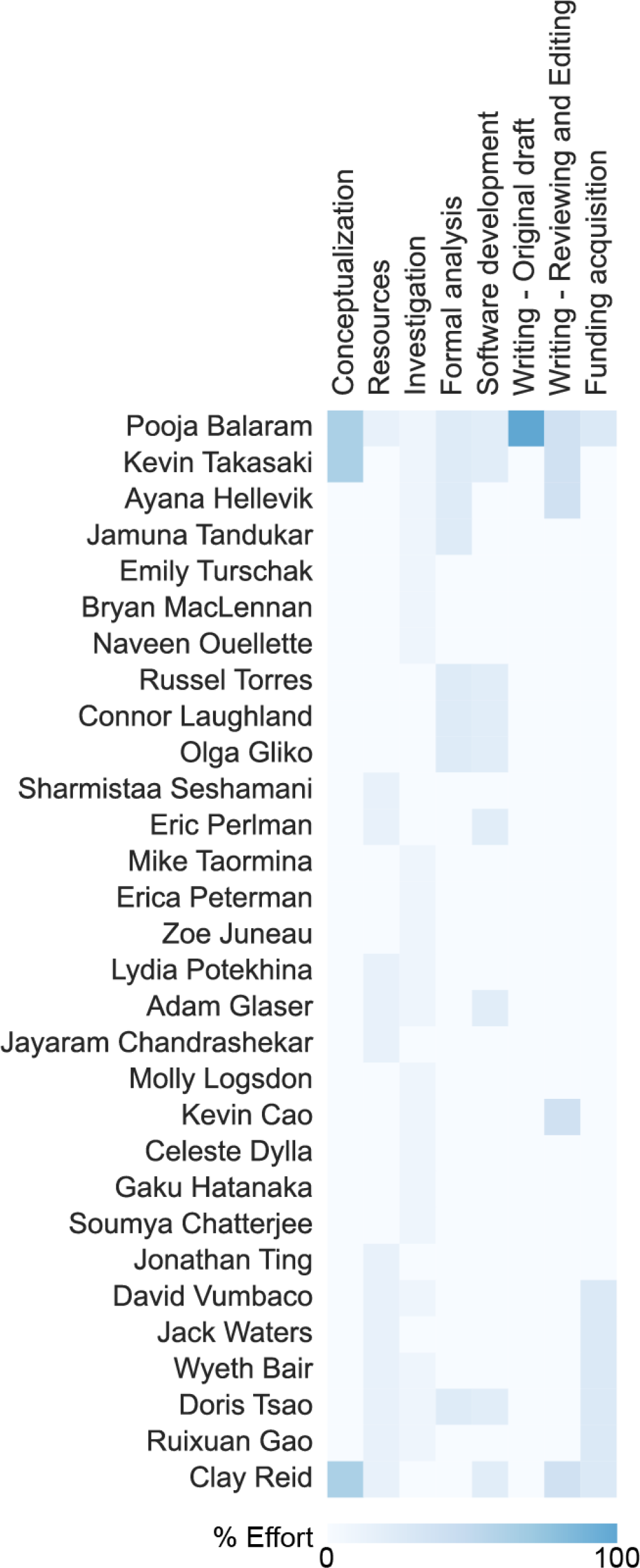

**Supplementary figure 1.**
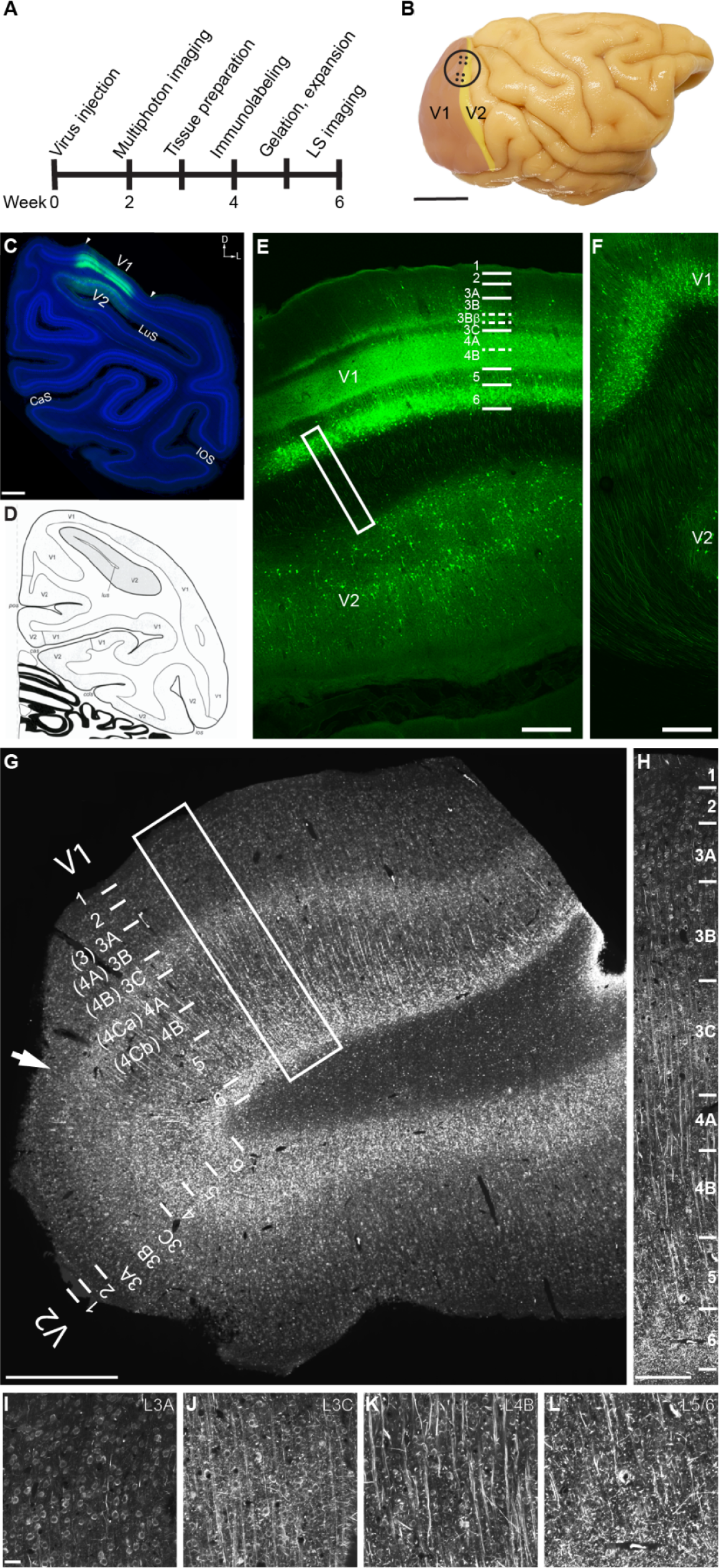
Combining optical physiology with expansion microscopy enables visualization of cellular morphology in functionally characterized neural circuits. (A) Experimental timeline. (B) Stereotaxic pressure injections (black dots) of PHP.eB-CAG-GCAMP6s or PHP.eB-hSyn-GCaMP6s were delivered along the V1/V2 border spanning the dorsal lunate gyrus prior to implantation of a cranial window (black circle) for multiphoton imaging. Scale bar is 1 cm. (C, D) Light microscopic imaging of coronal sections through injected craniotomy regions show extensive GCaMP6s expression across V1 and some expression in V2. Cortical areal boundaries are identified by laminar variations in DAPI labeling and comparisons with published atlases ^38^. Scale bars are 2 mm. (E, F) Confocal imaging of injected craniotomy regions reveals GCaMP6s-positive cell bodies in deep layers of V1 and V2 and putative axonal projections between V1 and V2, but similar patterns of expression in superficial cortical layers appear weak or nonexistent. White outline in E depicts the location of data in D. Scale bars are 500 *μ*m and 250 *μ*m respectively. (G-L) Following clearing, expansion and immunolabeling of SMI32, GCaMP6s-expressing cell bodies are clearly visible in all layers of V1 alongside SMI32-positive neurofilaments. White outline in G depicts the location of data in H, laminar boundaries within V1 are delineated based on SMI32. Scale bars are G:2 mm, H:150 *μ*m, I-L:50 *μ*m.

**Supplementary figure 2.**
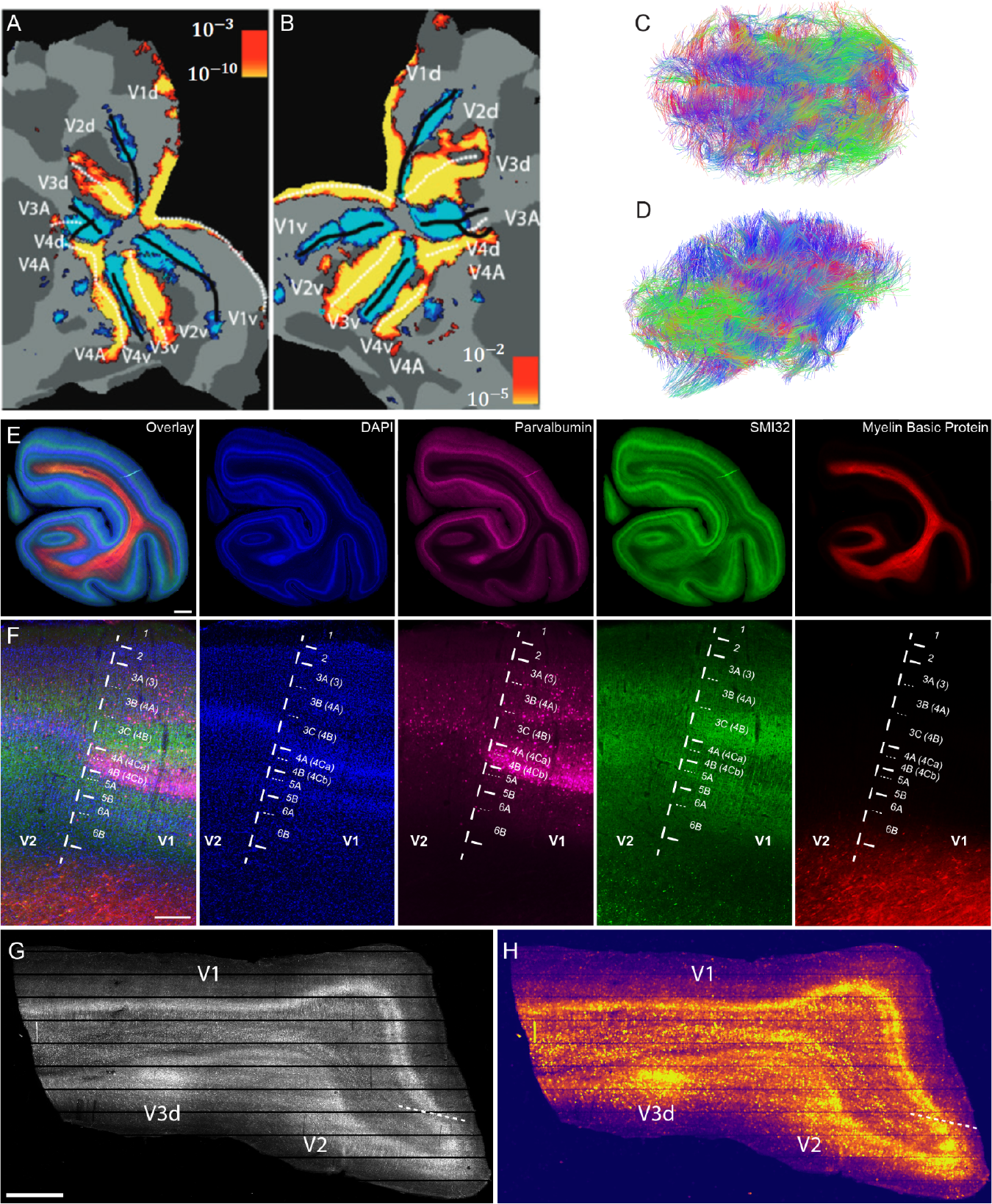
Combining magnetic resonance imaging with expansion microscopy facilitates the development of multi-scale multimodal proteomic atlases in large brains. (A,B) Maps of retinotopy spanning early visual cortical areas in one macaque monkey were initially acquired with in vivo 3T fMRI. Reproduced with permission from Hesse and Tsao, 2023. (C,D) Maps of connectivity were then acquired with 3T dMRI prior to tissue extraction and expansion microscopy. (E,F) Individual sections were fluorescently labeled to identify four common features of neural projections; cell nuclei (DAPI), myelin basic protein (MBP), non-phosphorylated heavy-chain neurofilaments (SMI32), and parvalbumin (PV). Cortical layers and areal boundaries can be demarcated based on variations in fluorescent label intensity and unique distributions of each cellular feature are clearly visible in each section. Scale bars are 1 mm and 100 *μ*m respectively. (G, H) A subset of sections were then gelled, cleared and expanded to visualize microscale distributions of each cellular feature. Exemplar hydrogel spanning the dorsal lunate V1/V2 border demonstrates clear demarcation of cortical areas and layers based on variations in fluorescent intensity.

